# Bottled aqua incognita: Microbiota assembly and dissolved organic matter diversity in natural mineral waters

**DOI:** 10.1101/154732

**Authors:** Celine C. Lesaulnier, Craig W. Herbold, Claus Pelikan, David Berry, Cédric Gérard, Xavier Le Coz, Sophie Gagnot, Jutta Niggemann, Thorsten Dittmar, Gabriel A. Singer, Alexander Loy

**Author notes:** Authors contributed equally to this study (Celine C. Lesaulnier, Craig W. Herbold, Gabriel A. Singer). Corresponding author: Alexander Loy, University of Vienna, Research Network Chemistry meets Microbiology, Department of Microbiology and Ecosystem Science, Division of Microbial Ecology, Althanstrasse 14, A-1090 Vienna, Austria.

## Abstract

**Background:** Non-carbonated natural mineral waters contain microorganisms that regularly grow after bottling despite low concentrations of dissolved organic matter (DOM). Yet, the compositions of bottled water microbiota and organic substrates that fuel microbial activity, and how both change after bottling, are still largely unknown.

**Results:** We performed a multifaceted analysis of microbiota and DOM diversity in twelve natural mineral waters from six European countries. 16S rRNA gene-based analyses showed that less than ten species-level operational taxonomic units (OTUs) dominated the bacterial communities in the water phase and associated with the bottle wall after a short phase of post-bottling growth. Members of the betaproteobacterial genera *Curvibacter*, *Aquabacterium*, and *Polaromonas* (*Comamonadaceae*) grew in most waters and represent ubiquitous, mesophilic, heterotrophic aerobes in bottled waters. Ultrahigh-resolution mass spectrometry of DOM in bottled waters and their corresponding source waters identified thousands of molecular formulae characteristic of mostly refractory, soil-derived DOM.

**Conclusions:** The bottle environment, including source water physicochemistry, selected for growth of a similar low-diversity microbiota across various bottled waters. Relative abundance changes of hundreds of multi-carbon molecules were related to growth of less than ten abundant OTUs. We thus speculate that individual bacteria cope with oligotrophic conditions by simultaneously consuming diverse DOM molecules.

## BACKGROUND

Bottled water, including natural mineral water, is an increasingly popular source of drinking water around the world and represents a multi-billion-dollar industry. The European Union regulates the exploitation and marketing of natural mineral waters to protect their unique characteristics and original purity. The latest EU Directive 2009/54/EC defines natural mineral water as microbiologically wholesome water of underground origin that is protected from all risk of pollution and can be clearly distinguished from other types of drinking water e.g. by its characteristic content of minerals and trace elements. Furthermore, disinfection or chemical treatment of natural mineral water is not permitted, yet it is routinely tested for its number of cultivable bacteria including several marker organisms (*Escherichia coli* and other coliforms, faecal streptococci, *Pseudomonas aeruginosa*, and sporulated sulfite-reducing anaerobes). The largely untreated nature of natural mineral waters allows that microorganisms from the source water aquifer and possibly the bottling plant (i.e. pipelines, storage tanks) act as inoculum for the bottle environment. Within a few days after bottling members of this ‘seed microbiota’ begin to grow during storage of natural mineral waters at ambient temperature, with absolute cell counts reaching 10^5^-10^6^ cells/mL [1-3].

Such microbial growth in non-carbonated bottled waters is a well-known fact, but the composition of the bottled water microbiota and its post-bottling dynamics have thus far been investigated by molecular techniques in only two natural mineral waters [2, 4]. These and numerous isolation-dependent studies [5-8] have established that *Alpha*-, *Beta*-, and *Gammaproteobacteria* are the prevalent microorganisms in bottled water. Beyond this, many fundamental questions regarding the microbial ecology of bottled waters still remain poorly answered. How different is the microbiota in bottled waters from different sources? How does the bottled water microbiota assemble? Are there differences between the free-living community in the water phase (plankton microbiota) and the inner-bottle-surface-associated community (biofilm microbiota)?

A persistent question that has puzzled researchers for decades is what substrates in bottled water may fuel the observed sudden microbial growth [8]? Although autotrophic growth has been suggested [4], it is commonly assumed that bottled water microorganisms mainly generate energy and multiply through heterotrophic utilization of dissolved organic matter (DOM) available in the bottle environment [8]. Ground water, treated drinking water, and oligotrophic surface freshwaters generally have low DOM concentrations in the range of 0.5-5 mg carbon/L. Microorganisms metabolize only a minor fraction (0.01-0.1 mg carbon/L) of this DOM pool while a larger refractory fraction remains untouched [9]. The susceptibility of DOM to microbial turnover depends on its origin and diagenetic history, which define chemical composition [10, 11], and its concentration, which defines the probability of microbial encounter for a specific chemical structure [12]. DOM analytics have been dramatically advanced by the advent of Fourier transform ion cyclotron resonance mass spectrometry (FT-ICR-MS) with ultra-high mass resolution allowing to accurately identify thousands of individual molecular formulae in a single environmental sample [13]. While FT-ICR-MS has provided fundamental insights into the molecular diversity of DOM and its turnover in various aquatic environments such as oceans [12-14], lakes [15], glaciers [11], and groundwater [16, 17], analogous information is not available for bottled drinking water.

Here, we simultaneously analyzed microbiota and DOM composition in twelve European non-carbonated natural mineral waters and their corresponding source waters by multiplexed sequencing of bacterial 16S rRNA gene amplicons, FT-ICR-MS, and complementary analysis of microbiota and physicochemical parameters. Our approach included fine-scale temporal investigation of two representative bottled waters over two months of incubation and detailed comparative analysis of all twelve bottled waters one day after filling in polyethylene terephthalate (PET) bottles. We revealed corresponding patterns in DOM turnover and microbial community development after bottling and thereby provide novel insights into the molecular microbial and chemical ecology of this important drinking water source.

## METHODS

### Sampling and storage of natural mineral waters

We analyzed non-carbonated natural mineral waters retrieved from twelve European bottling plants of well-derived mineral water in the years 2011 and 2012 (Supplementary Tables S1-S4). Samples from each plant included bottled water and corresponding well water; the latter was taken before its entry into the bottling plant from a sampling port located at the head of each source. Bottled water 6 receives water from two wells 6a and 6b, while bottled waters 9a and 9b receive water from the same well 9. Bottled and well waters were filled in brand-specific 0.5 L and standard 1 L PET bottles, respectively. Water bottles were transported to the laboratory within 24 hours, and were subsequently stored in a dark, climate-controlled room (20-24°C) for sampling at regular time points after bottling. All bottled waters used in this study complied with the legal microbiological criteria (EU Directive 2009/54/EC).

### Recovery of plankton and biofilm biomass

All waters were sampled in triplicate at each time point. Each replicate sample consisted of biomass from one or more separate bottles of water. Planktonic microorganisms were recovered on polycarbonate filters with a pore size of 0.22 μm (GTTP04700; Millipore, Eschborn, Germany) using a stainless steel vacuum filtration unit equipped with three 500 mL filter funnels (Sartorius, Göttingen, Germany). Per replicate sample, 3 L (e.g. 6x 0.5 L bottles or 3x 1 L bottles) of water were filtered for nucleic acid extraction and 0.5 to 3 L for microscopy. Filters with cellular biomass were air dried, cut in half, and stored at -80°C for DNA extraction or fixed with para-formaldehyde and stored at -20°C for microscopy [2]. Biofilm samples were recovered from the inner linings of the same bottles that were used for harvesting planktonic biomass. Bottles were cut in half with a sterile scalpel blade and the entire inner surface of each bottle was swabbed with a sterile viscose collection swab (Deltalab, Carcassonne, France). The swab tips were then cut off and stored at -80°C prior to DNA extraction.

### Quantitative fluorescence microscopy

FISH and DAPI-staining of microbial cells on polycarbonate filters was carried out as described previously [2]. Fluorescent cells were quantified by analyzing 20 pictures of randomly chosen fields of view per replicate sample with the image analysis program DAIME [18]. The following fluorescently-labelled probes were applied for FISH using stringent hybridization conditions and, if applicable, together with their unlabeled competitor probes: EUB338 probe mix for *Bacteria*, ARCH915 for *Archaea*, ALF968 for *Alphaproteobacteria*, BET42a for *Betaproteobacteria* (GAM42a as competitor), AQUA827 for genus *Aquabacterium*, MEVE845 for the genus *Methyloversatilis*, and HGC69a for *Actinobacteria* [19]. Fluorescently-labeled NONEUB probe was applied as negative control to all analyzed samples in order to assess unspecific-binding. Probe MEVE845 (S-G-MEVE-0845-a-A-21, 5’-TTA GCT GCG GTA CTC AAT GAG-3’) and a corresponding competitor probe c1MEVE845 (5’-TTA GCT GCG TTA CTC AAT GAG-3’) were newly designed based on the non-redundant ARB-SILVA database version 111Ref [20] using the probe tools of ARB [21], probeCheck [22], and RDP II [23]. Dissociation-curve analysis [18] with the perfectly-matched type strain *Methyloversatilis universalis* showed that probe MEVE845 should be applied at 25% formamide to ensure its specificity.

### DNA extraction

Polycarbonate filters with planktonic biomass were first cut into smaller pieces with sterile scissors. Each sample (i.e., filter with planktonic biomass or swab tip with biofilm biomass) was transferred into an individual 2 ml Lysis A matrix tube without the 1/4 inch ceramic sphere (MP Biomedicals LLC, Solon, OH, USA). Cell lysis was initiated by adding 400 μl of lysis buffer (10 mM Tris-HCl, 1 mM EDTA, 100 mM NaCl, 0.5% SDS, 20 μg proteinase K, pH 8) and incubating at room temperature for 15 minutes. Subsequently, 500 μl of phenol-chloroform-isoamylalcohol (25:24:1, Carl Roth Karlsruhe, Germany) was added, followed by vortexing for 2 minutes at maximum speed and centrifugation at 13,000 rpm for 2 minutes. The supernatant was recovered and further purified by a second round of phenol-chloroform-isoamylalcohol treatment. DNA in the recovered supernatant was precipitated by adding 2.5 volumes of ice-cold 100% ethanol, 1 μl 3 M sodium acetate (pH 5) and 1 μl glycogen and incubation at -20°C for 2 hours. Precipitated DNA was recovered by centrifugation at 13,000 rpm for 10 minutes and washed with ice-cold 70% ethanol. Purified DNA was air dried, dissolved in 50 μl sterile TE buffer (10 mM Tris-HCl, 1 mM EDTA, pH 8) and stored at -20°C.

### Quantification of nucleic acids

DNA extracts and PCR products analyzed by agarose gel electrophoresis and quantified by using the Quant-iT PicoGreen dsDNA Assay Kit (Invitrogen Corporation, Carlsbad, CA, USA). Samples and DNA standards were prepared as per manufacturer’s instructions. Samples were placed in a black, flat bottom, 96 well plate (Greiner bio-one, Frickenhausen, Germany) and analyzed with a Microplate Reader (Tecan Infinite M200; Tecan Group Ltd, Männedorf, Switzerland).

### Multiplex amplicon pyrosequencing and sequence analysis

Barcoded 16S rRNA gene amplicons for 454 pyrosequencing were prepared from planktonic and biofilm DNA using a previously published 2-step PCR procedure and primers (909F, 5′-ACTCAAAKGAATWGACGG-3′ and 1492R, 5′-NTACCTTGTTACGACT-3′) that amplify variable regions V6 to V9 of the 16S rRNA gene of most bacteria [24]. To account for potential contamination [25], PCRs without addition of a DNA template or with DNA extracts from empty swap tips and empty polycarbonate filters were performed during each PCR run. Most negative control PCRs did not yield a visible amplicon, except of six that showed a faint band in the agarose gel and were thus also sequenced. Pyrosequencing of barcoded amplicon pools was performed with Titanium reagents on a 454 genome sequencer FLX (Roche, Basel, Switzerland) by the Norwegian Sequencing Centre (Oslo, Norway) or by Eurofins Genomics (Ebersberg, Germany).

Raw 454 pyrosequencing flowgrams were denoised and checked for chimeras using AmpliconNoise [26] within the QIIME environment [27]. Denoised reads were subsequently clustered into OTUs at 97% identity, roughly corresponding to species-level OTUs [28], with UPARSE and an OTU table was generated with the associated <u>uc2otu.py</u> script [29]. Taxonomic classifications were assigned using the Ribosomal Database Project naïve Bayesian classifier [30] using the Silva SSURef database v119 [31]. The OTU table and taxonomic classifications were imported into the R environment [32] for all further analysis. Contaminant OTUs were defined as OTUs that had higher relative abundance in negative controls than in samples. Contaminant OTUs and OTUs observed with fewer than three total counts across all samples were removed for all subsequent analysis. After sequence quality, contamination, and chimera filtering, a total of 567,310 reads from 215 individual samples, i.e. 2639 ± 1650 (mean ± SD) reads per sample, were retained (Supplementary Table S5).

Shannon and Simpson metrics were calculated using diversity() and estimateR() functions from the vegan package [33]. Goods coverage was calculated as 1 – (number of singletons/number of reads) for each sample. Weighted UniFrac distance metrics were calculated using the UniFrac function from the Phyloseq package [34] and principal coordinate plots (PCoA) plots were generated using the rda() function in the vegan package. Specific hypothesis tests reported in the main text were conducted using cor.test(), wilcox.test() and/or t.test() from the statistics package in R. Multiple testing was accounted for using the Benjamini-Hochberg procedure with the p.adjust(method=”fdr”) function. For all hypothesis tests and PCoA plots, the OTU table was re-sampled at 500 reads per sample and any sample with fewer than 500 reads was omitted. This procedure was repeated 100 times and mean p-values reported for hypothesis tests. Multiple re-samplings were visualized in PCoA plots by fitting sample coordinates together using the procrustes() function from the vegan package. Heatmaps show only OTUs that exceed 1% relative abundance in relevant samples and associated dendrograms were calculated using the hclust(method=”average”) function using distances calculated using dist.dna(pairwise.deletion=TRUE) on an alignment of reads constructed with mafft -linsi [35].

‘Growing OTUs’ were determined by correlating estimated OTU cell numbers to the number of days after bottling in a resampling scheme. First, total cell numbers for a sample were estimated using the rnorm() function with the measured mean and standard deviation as the input distribution. Then, samples were resampled as previous but used to calculate a resampled relative abundance. Resampled relative abundance was subsequently multiplied by estimated DAPI-cell count to estimate the number of cells from each OTU in a resampled community. During each resampling iteration, Pearson correlations and associated p-values were calculated between log(cell count+1) and log(day after bottling). The resampling procedure was repeated 100 times and the median p-values for each OTU was adjusted using the Benjamini-Hochberg procedure with the p.adjust(method=”fdr”) function. ‘Growing OTUs’ were defined as those OTUs with a significant adjusted median p-value (<0.05) for a positive correlation of estimated cell number with time.

Three sets of ‘core OTUs’ were defined as those seen in (1) a majority of well waters, (2) a majority of early bottled waters (1 day after bottling) or (3) a majority of late bottled waters (≥14 days after bottling). In each case “majority” is in regard to the number of distinct water types (n=12), not samples. A final set of ubiquitous OTUs was defined as those OTUs present in all three core OTU definitions.

### Clone library preparation and Sanger sequencing

Almost full-length bacterial 16S rRNA genes were amplified with primers 616V (5’-AGAGTTTGATYMTGGCTC-3’; [36]) and 1492R (5’-NTACCTTGTTACGACT-3’; [24]). Individual PCRs were carried out using planktonic DNA samples from Water 1 (3 and 28 days after bottling) and Water 2 (6 and 28 days after bottling) obtained in the year 2011, the Taq DNA polymerase kit (Fermentas Inc., Hanover, MD, USA), and nucleotide-mix (2 mM/dNTP) (Fermentas Inc., Hanover, MD, USA) as per manufacturer’s instructions and at an annealing temperature of 51°C and with 25 cycles. PCR amplicons were cloned using the pCR2.1 TOPO TA cloning kit (Invitrogen Corporation, Carlsbad, CA, USA) as per manufacturer’s instructions. Plasmid DNA from each clone was purified using the QuickLyse Miniprep Kit (Qiagen, Venlo, The Netherlands) and inserts were sequenced by Microsynth (Balgach, Switzerland). The recovered sequences were manually proofread using the sequence and chromatogram software FinchTV V1_4_0 (Geospiza/Perkin Elmer, Seattle, WA, USA).

### Phylogenetic Analyses

Near full-length 16S rRNA gene sequences and representative reads of 454-derived OTUs that were identified as core and/or growing OTUs were used for phylogenetic reconstruction. Appropriate reference sequences were identified using a combination of blastn, the Ribosomal Database Project sequence match tool and the ARB software package with the Living Tree Project database release 108 [21, 37]. In cases in which the 454-derived OTU sequence completely matched a full-length 16S rRNA clone or reference sequence, only the full-length sequence was used for treeing. The final set of sequences was aligned using SINA [38] and the phylogeny was calculated using both RAxML [39] and Phylobayes 3 [40]. The RAxML tree was calculated using the GTRGAMMA model with 1000 rapid bootstraps. The Phylobayes tree was calculated using the GTR model with 4 gamma-distributed rate categories and 5 independent chains of 40,000 generations. The first 10,000 generations of each chain were discarded as burn-in for posterior probability calculations.

### Physicochemical analyses

Prior to biomass recovery, the oxygen concentration in the water bottles was measured using a needle-type oxygen microsensor and a Microx TX3 Micro fiber optic oxygen transmitter (PreSens Precision sensing, Regensburg, Germany). Calibration and measurements were performed as per manufacturer’s instructions. PET bottles were pre-pierced with a sterile needle followed by immediate injection of the microsensor optode into the water for oxygen measurements. Additional water parameters were measured by the Nestlé Quality Assurance Center using potentiometry (e.g. pH, conductivity, alkalimetry), Flow Injection Analysis (e.g. nitrate, nitrite, ammonium, orthophosphate), inductively coupled plasma mass spectrometry (e.g. e.g. calcium, magnesium, sodium, potassium, sulfate, silica, iron, manganese).

### Assimilable organic carbon

The amount of organic carbon that was assimilated by the growing microbial community was inferred based on total cell counts at the beginning and end of microbial growth as previously described [41].

### Dissolved organic matter extraction

DOM from all samples was extracted from respective volumes (0.5-5 L) corresponding to approximately 0.25-1 mg total dissolved organic carbon. Water was filtered directly from the original, brand-specific PET bottles (0.5 L) or standard PET sampling bottles (well water) through a double layer of pre-combusted (450°C, 4 h) glass fiber filters (0.7 μm pore size, Whatman GF/F) into acid-washed pre-combusted glassware. All bottled water and well water samples were analyzed in triplicate per time point; for each replicate water from independent original product or sampling bottles (0.5-1 L) was pooled as necessary to reach adequate volumes. In parallel to each set of samples, we filtered equal volumes of Milli-Q water as blank DOM controls. Filtered water was then used for extraction of DOM [42] on a solid phase (Agilent Bond Elut PPL 3ml 100 mg cartridges, VWR, Arlington Heights, USA) after acidification to pH 2 (Suprapur-grade HCl, Carl Roth, Mannheim, Germany). DOM was eluted from cartridges with LC-MS-grade methanol (Sigma Aldrich, St Louis, USA) and stored in pre-combusted 4 mL amber glass vials at -20°C pending FT-ICR-MS. We quantified dissolved organic carbon in the original filtered water by wet-chemical oxidation (Sievers 900 TOC Analyzer operated with an inorganic carbon removal unit). We determined a method detection limit of the Sievers 900 according to US EPA guidelines of approximately 6 μg C L^-^1.

### FT-ICR-MS and MS-data preprocessing

Mass spectrometry of DOM extracts (adjusted to 20 ppm carbon in 1:1 methanol/ultrapure water) was done on a 15 Tesla Solarix FT-ICR-MS (Bruker Daltonics, Bremen, Germany) in electrospray ionization (ESI) negative mode (500 accumulated scans, 2 sec ion accumulation time) searching for masses from 153 to 2000 Da. No peaks were detected for masses >1000 Da. Following internal calibration, peaks with S/N>3 were exported from Bruker-DataAnalysis software for further data analysis using in-house code in R [32] (Supplementary Table S8). A first assignment of molecular formulae to peaks was done assuming single-charged deprotonated molecular ions and Cl-adducts for a maximum elemental combination of C_100_H_250_O_80_N_4_P_2_S_2_, with a mass tolerance of 0.6 ppm, and using the following restrictions: agreement with the nitrogen rule, positive integer double bond equivalent for uncharged molecule, minimum C_1_H_1_O_1_, P<(O+1), S<(O+1), H:C within [0.3, 2.5], O:C and N:C within [0,1], H≤2C+2+N, at least 1 O for each P or S, and no heteroelement (N, S, P) co-occurrence. We then checked for isotope confirmation of all potentially valid formulae using generated isotope intensity patterns (up to 10 daughter peaks considering isotopes of all elements except P) and based on adequate mass shift(s) (tolerance 0.6 ppm) and adequate intensity ratio(s) (±40%) of isotopic daughter peaks to the monoisotopic, parent peak [43]. For formulae assigned to the molecular groups of condensed polyaromatics, saturated fatty acids and carbohydrates and involving heteroatoms (N, S, P) (see below), more stringent limits were set for isotope confirmation (halved tolerance and halved maximum deviation from the expected peak ratio). A single daughter isotope peak sufficed for confirmation of a suggested sum formula, 2 daughter peaks were minimum for sum formulae with Cl, which has abundant secondary isotopes and produces prominent daughter peaks besides those produced by exchange of ^12^C by ^13^C. In case of multiple assignments to the same peak, we gave preference to (i) extremely abundant molecules commonly found in environmental samples (CHON, CHON_2_, CHOS), (ii) formulae with better isotope confirmation (more daughter peaks and isotope confirmation across a greater number of samples), and (iii) formulae involved in longer homologous series based on CH_2_ (aliphatic elongation) and CO_2_ (acid-based elongation). Intensities of formulae found in deprotonated charged state and as a Cl-adduct were summed. Finally, we aligned detected masses across samples based on assigned formulae, keeping only formulae that achieved stable isotope confirmation in at least one sample [11]. 126 formulae assigned to mass peaks with S:N>5 found in any of a total of 20 blank samples were considered to be contaminants and deleted from the dataset.

### DOM data analysis

FT-ICR-MS data is graphically presented in van Krevelen plots, which show identified sum formulae in a space defined by O:C (oxygen richness) and H:C (saturation) ratios; plotting order was random to avoid bias created by systematic overplotting as is common in van Krevelen plots showing thousands of compounds. Before statistical analysis, we further filtered the dataset based on a ‘replicate filter’, i.e., any singlet molecular formula determined for a set of replicates was deleted [44]. As replicate sample sets we considered all samples from the same source and incubation time. This filter was not applied to the single replicate of Well 8. The final dataset consisted of 4055 sum formulae. To condense these information, we grouped molecular formulae into 12 non-overlapping molecular groups based on elemental composition and derived structural information such as double bond equivalents (DBE) and a computed aromaticity index [45]; the most prevalent categories are reported and defined in Supplementary Table S7. While this categorization is to some degree arbitrary, it allows an overview of the molecular data. As descriptors of molecular diversity, we report the total number of identified sum formulae (richness), the Shannon-Wiener index and evenness. To investigate compositional changes of DOM over time or due to geographical variation among wells, we used principal component analysis (PCA) based on centered log-transformed relative intensity data (separately for the ‘time-course experiment’ datasets of Water 1 and Water 2, and for the ‘diversity study’ dataset). Time course data was also used in redundancy analysis [46] with incubation time as the single constraint to identify significance of temporal changes of DOM composition. Principal components, i.e., gradients of major compositional variation of DOM, were (i) plotted against other variables (e.g., incubation time), and (ii) correlated (Spearman) with relative intensities of individual molecular formulae for color-coding molecules in the van Krevelen space. For the time series datasets, we also opposed sets of molecular formulae with very low/high correlation coefficients (<20% and >80% quantiles) with respect to their location in van Krevelen space, dominance of molecular groups and molecule mass. These two formula sets serve to describe two pools of compounds likely decreasing and increasing during incubation; it is impossible to unequivocally identify decrease or increase of a compound from relative intensity data. All data analysis was carried out in R version 3.2.1 [32] using the packages vegan [33] and MASS [47].

## RESULTS

### Bacterial community and dissolved organic matter composition during microbial growth in two representative bottled waters

We analyzed microbial growth, physicochemical properties, bacterial community composition, and molecular DOM composition in two non-carbonated bottled waters, Water 1 from France and Water 2 from Poland, during 56 days of storage after filling in brand-specific PET bottles at the bottling plant. The two waters were selected to represent the range of DOM concentrations (Water 1: <0.06 mg carbon/L, Water 2: 1.0-1.2 mg carbon/L) that are typical for bottled waters at the time of bottling (Table 1, Supplementary Tables S1 to 4). This ‘time-course experiment’ was performed twice in two consecutive years (2011, 2012), but in-depth analysis of DOM was restricted to 2012 when we also conducted an analysis of the well waters sampled into identical standard PET bottles at the bottling plant. Generally, small temporal variations in generic physicochemical parameters were observed in both waters during storage (Table 1, Supplementary Tables S1 and S2). The two waters differed in mineral content and with respect to concentrations of dissolved oxygen, the latter is likely because of the differences in initial pressure under which these waters were bottled (Table 1, Figure 1). Microorganisms were not limited by oxygen availability in either water as oxygen concentrations were never below 6.9 mg/L (Figure 1) [48]. Microorganisms multiplied in both waters as observed for other non-carbonated bottled waters [2]. Within a week, total cell counts increased several fold to ≥10^5^ cells/mL and remained at this concentration until the end of the experiment. While Water 2 contained >20 times more DOM than Water 1 (Table 1), only about 2-4 times more organic carbon was assimilated by the microbiota in Water 2 (18.2 μg/L in year 2011, 40.1 μg/L in year 2012) than Water 1 (10.4 μg/L in year 2011, 20.9 μg/L in year 2012). This indicated that only a small fraction of total DOM was immediately available for microbial growth, as has been shown for other types of oligotrophic drinking waters [9].

**Figure 1.**
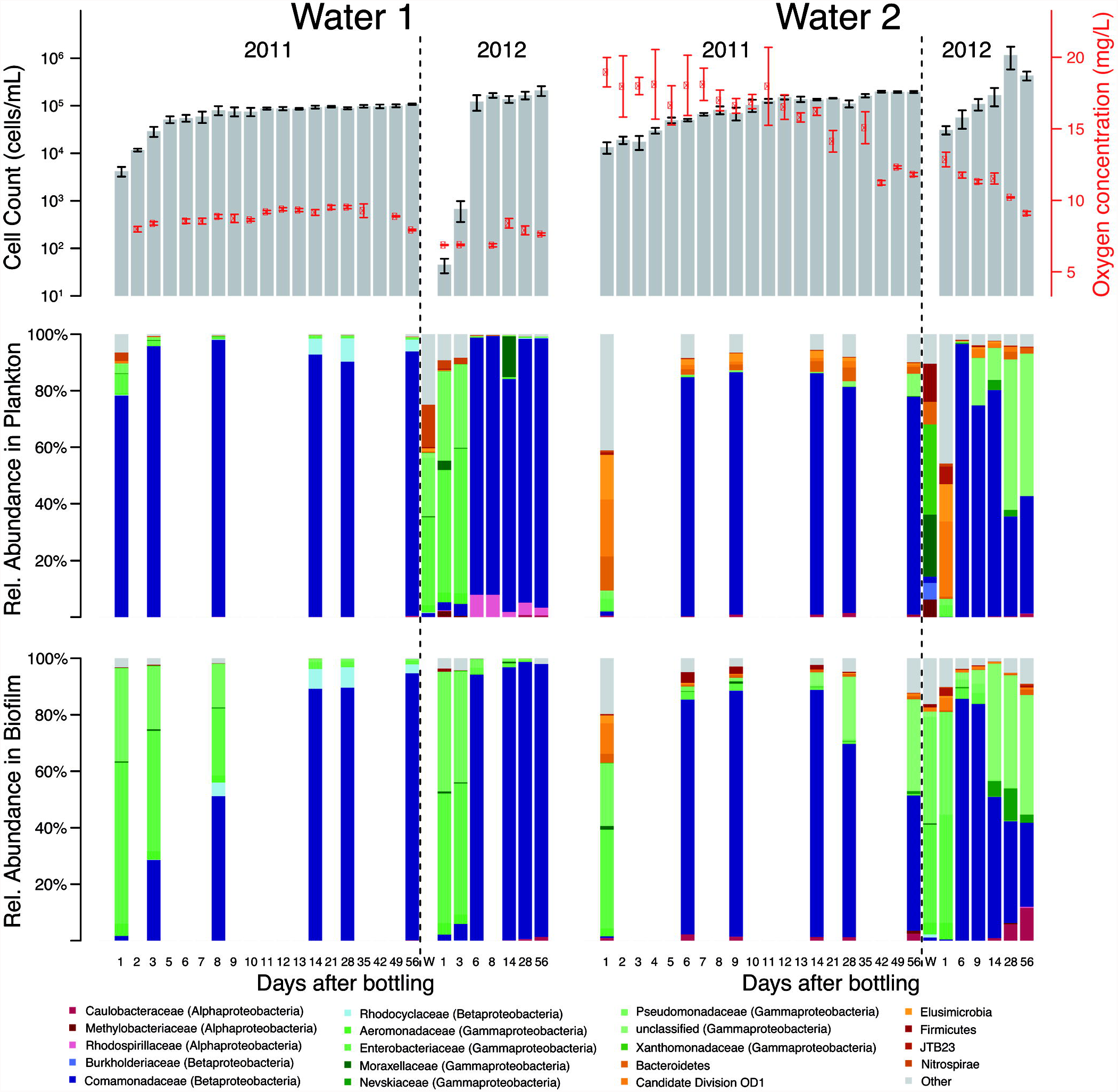
Microbial growth, bacterial community shifts, and oxygen concentration in non-carbonated natural mineral water 1 and 2 after bottling. In years 2011 and 2012, Water 1 and 2 were filled in PET bottles and monitored during 56 days of storage. Total planktonic cell counts (grey bars) and dissolved oxygen concentrations (red) are shown as mean ± standard error (n = 3). To ensure bottle stability during packaging and delivery, bottling is performed using compressed air with an initial pressure of 1.5 bar for Water 1 and 2.2 bar for Water 2. W, well water. Temporal changes in bacterial community composition in the water (plankton) and on the inner bottle wall (biofilm) were analyzed by 16S rRNA gene amplicon sequencing. Bar charts only show phyla with >5% relative abundance in at least one sample. *Proteobacteria* are shown as individual families.

**Table 1.**
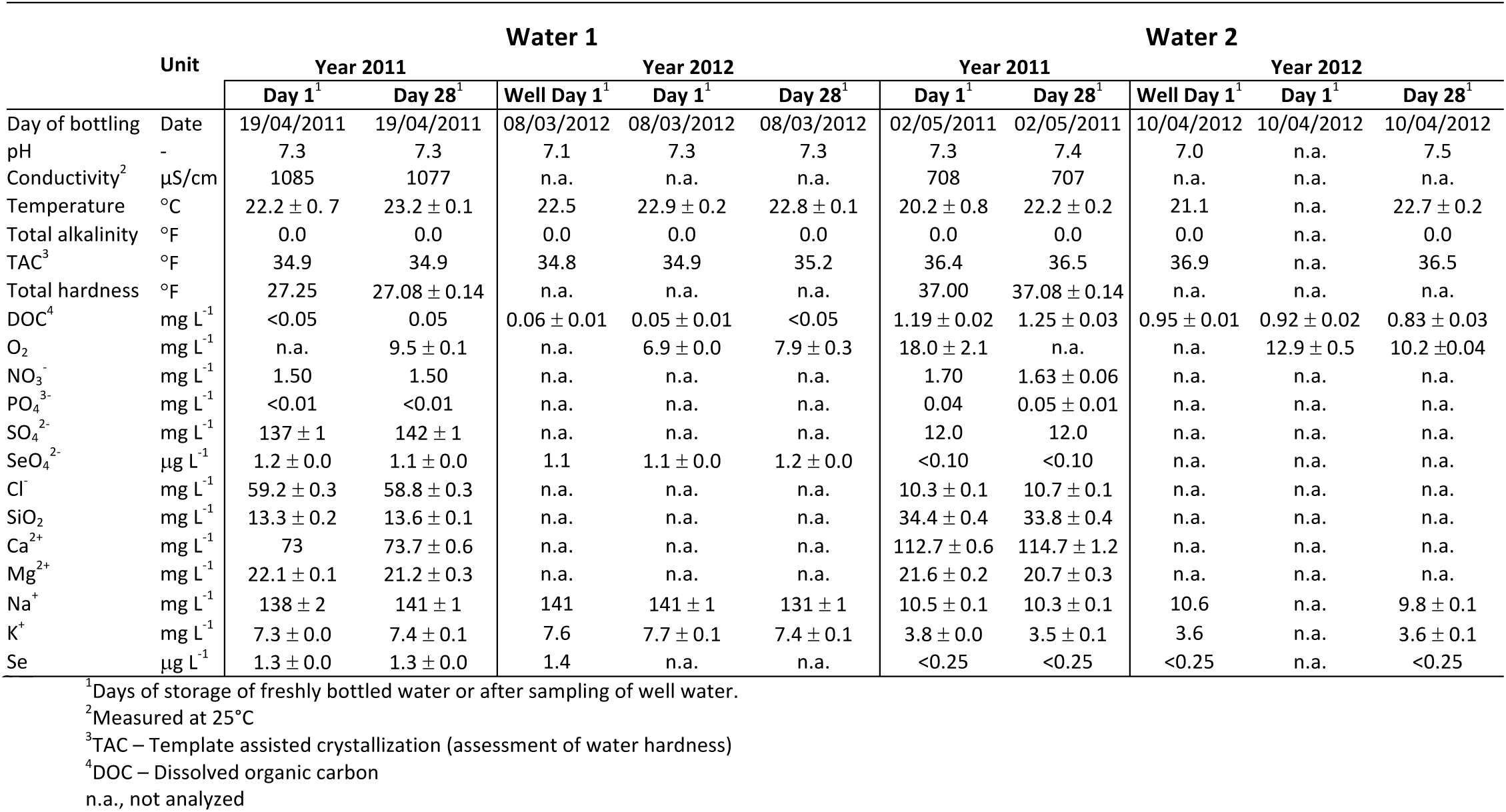
Physicochemical water properties and further sampling information from selected time points during time-course analyses of natural mineral waters 1 and 2 after bottling. See Supplementary Tables S1 and S2 for extended versions of this table with all sampling time points.

Initial screening of the two bottled waters during and after microbial growth by fluorescence in situ hybridization (FISH) with domain-specific probes showed a prevalence of bacteria (Supplementary Figure S1), which corroborated previous studies that archaea and eukaryotic microorganisms are absent or low in abundance in bottled groundwater [2, 4, 49]. We thus focused all subsequent microbial analyses on the bacterial community. Temporal variations in planktonic and biofilm community structure after bottling were revealed by multiplex pyrosequencing of bacterial 16S rRNA gene amplicons. The retrieved, quality-filtered sequences in the time-course experiment dataset consisted of 672 species-level operational taxonomic units (OTUs), with 127 having ≥1% relative abundance in at least one sample. 70 and 115 of these abundant OTUs were detected in Water 1 and Water 2, respectively, of which 58 OTUs were present in both waters and 10 grew over the time-course experiment (Supplementary Tables S5 and S6).

Bacterial diversity within each sample (alpha-diversity: Chao1, Shannon index, Simpson index) was overall higher in Water 2 than in Water 1 (Supplementary Table S5, median resampled p<1x10^−9^, two-sample t-test for all alpha diversity metrics). Onset of planktonic microbial growth in both waters was accompanied by a considerable decrease in both alpha-diversity (median resampled p<0.015, for correlation of any alpha diversity measure in either water against log(days after bottling)) and pair-wise diversities between all samples at a given time point (beta-diversity: median resampled p<0.024, for correlation of unifrac distance against log(days after bottling) for each water). Planktonic and biofilm communities within and between the two waters were clearly different early after bottling but converged into very similar communities within nine days of storage (Figures 1 and 2). After day 7, when most of the increase in total cell counts had occurred, only 1-6 (median 2) and 4-15 (median 6.5) OTUs made up ≥90% of the relative 16S rRNA gene read count throughout the remaining incubation time in Water 1 and 2, respectively.

**Figure 2.**
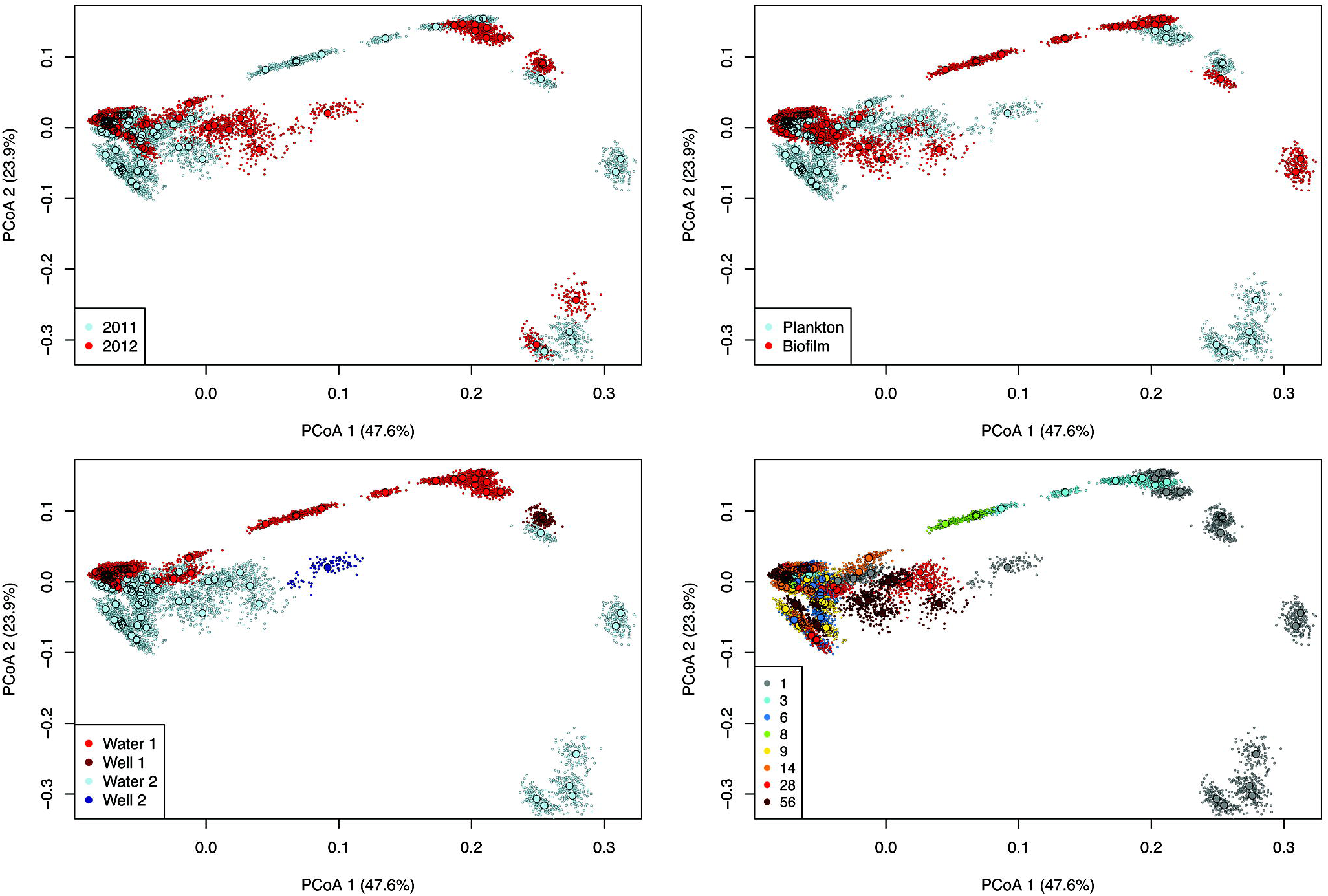
Microbiota beta-diversity in natural mineral water 1 and 2 after bottling. PCoA analysis based on weighted UniFrac distances calculated from bacterial 16S rRNA gene amplicon data. Each larger circle indicates the microbiota of an individual, replicate water sample. *In-silico* dataset re-sampling is visualized in PCoA plots as smaller circles. Each panel shows the same PCoA plot with individual coloring according to the year of sampling (year 2011 vs year 2012), type of microbiota (plankton vs biofilm), type of water (well water vs bottled water; Water 1 vs Water 2), and days after bottling (between 1 and 56 days).

In Water 1, 16S rRNA genes from *Gammaproteobacteria* (*Enterobacteriaceae*, *Pseudomondaceae*) generally dominated in low-biomass well water and bottled water samples from early time points (Figure 1). However, members of the class *Betaproteobacteria* (*Comamonadaceae*) contributed most strongly to microbial growth and continued to dominate the planktonic and biofilm communities throughout the incubations. Early after bottling, Water 2 had higher phylum/class/family-level diversities than Water 1, with considerable representation of 16S rRNA genes from the *Alpha*-, and *Gammaproteobacteria*, *Bacteroidetes*, *Elusimicrobia*, and the candidate phyla radiation lineage OD1 (*Parcubacteria*) (Figure 1) [50]. Microbial growth in Water 2 was also mainly due to *Betaproteobacteria* (*Comamonadaceae*).

Regarding species-level OTU composition immediately after bottling, the gammaproteobacterial OTUs 3, 4, and 21 were most abundant in 16S rRNA sequence libraries of both waters (Supplementary Figure S2). Subsequent microbial growth in both waters and in both years was attributed to OTUs 1 (*Curvibacter*, *Comamonadaceae*) and 2 (*Aquabacterium*, *Comamonadaceae*) (Figure 3, Supplementary Table S6). In addition to these two ubiquitous OTUs, there were further OTUs that grew inconsistently, usually only in a single year for a particular water source. For example, OTU 13 (*Methyloversatilis*, *Rhodocyclaceae*) grew only in Water 1 from 2011. Instead, OTUs 25 (unclassified *Xanthomonadales*), 68 (unclassified *Xanthobacteraceae*), and 1408 (unclassified *Comamonadaceae*) contributed to bacterial growth in Water 2 in 2011. Different OTUs contributed to growth in Water 2 samples from 2012, namely OTU 9 (unclassified *Xanthomonadales*) and OTU 993 (*Polaromonas*, *Comamonadaceae*). For phylogenetic analysis, we recovered near full-length bacterial 16S rRNA gene sequences from year-2011-samples of both waters at later post-bottling time points (Figure 3). Several OTUs had 100% identity to near full-length 16S rRNA sequences from this and other studies. For example, sequences of OTUs 1 and 2 perfectly matched *Curvibacter fontanus* AQ12 (AB120966) [51] and *Aquabacterium parvum* B6 (AF035052) [52], respectively. FISH experiments confirmed the dominance of *Betaproteobacteria* and the high relative abundances of specific genera during and after microbial growth in the respective water samples (Supplementary Figure S1).

**Figure 3.**
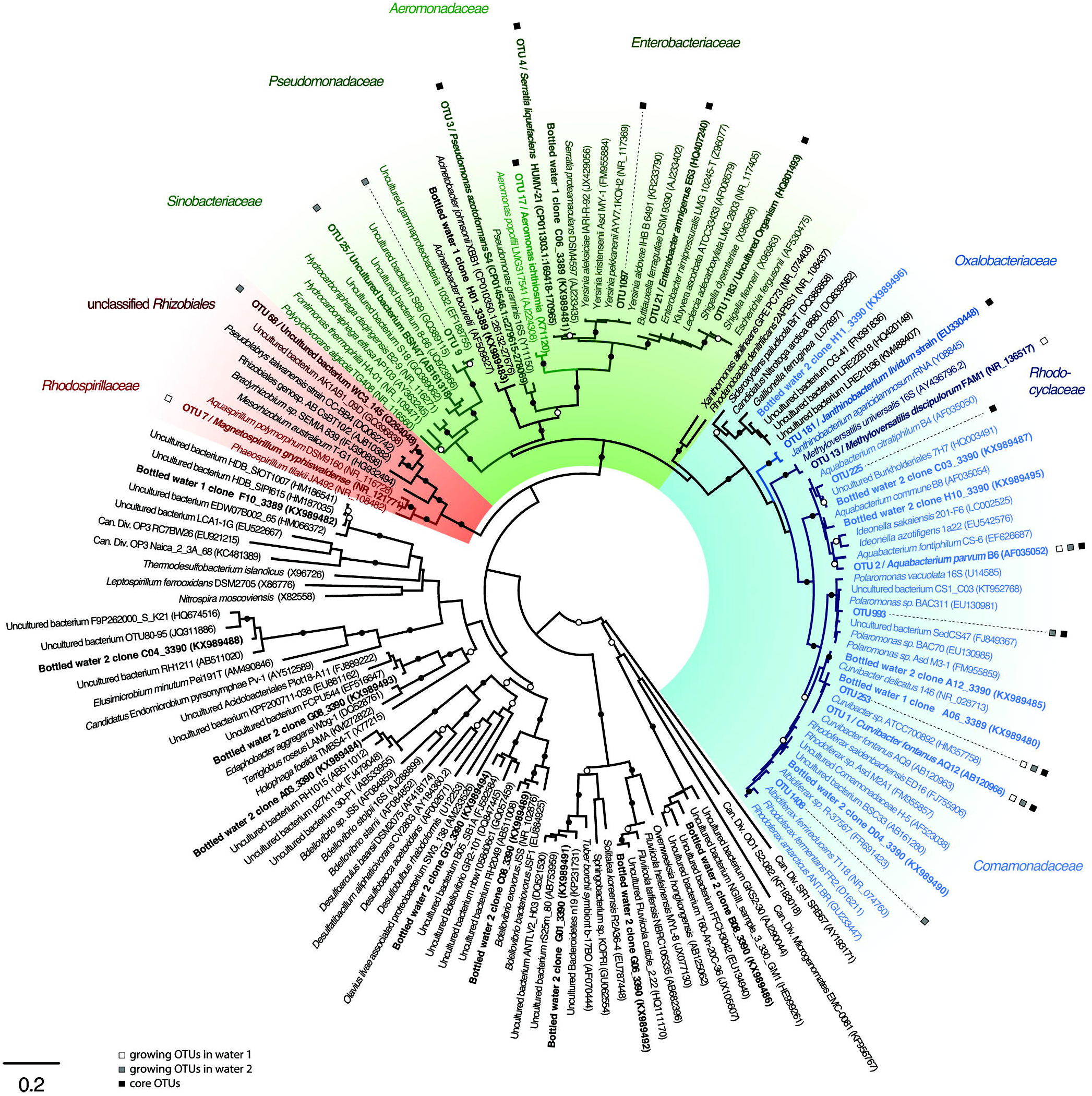
16S rRNA gene tree showing the affiliation of core and growing bottled water bacteria. Near full-length 16S rRNA gene sequences and representative reads of 454-derived OTUs that were identified as core and/or growing OTUs were used for phylogenetic reconstruction. Core and/or growing OTUs and near full-length sequences recovered in this study are shown in bold. White and grey squares indicate OTUs growing in Water 1 and Water 2, respectively. Black squares indicate core OTUs. The bar indicates 2% estimated sequence divergence. Closed circles indicate >95% RAxML bootstrap support and >0.99 Phylobayes posterior probability, respectively. Open circles indicate 70-95% RAxML bootstrap support and 0.90-0.99 Phylobayes posterior probability, respectively. *Alpha*-, *Beta*-, and *Gammaproteobacteria* are shaded in red, blue, and green, respectively.

FT-ICR-MS analysis of water samples taken during the time-course experiment in 2012 revealed high DOM complexity and similar temporal dynamics despite considerable differences in total DOM concentrations and composition between the two waters (Supplementary text) (Figures 4 and 5, Supplementary Table S3 and S4). We identified a total of 3152 and 3177 different molecular formulae across the time-course experiment of Water 1 and 2, respectively. Each water sample contained on average more than 2500 different molecular formulae (Water 1: 2496±135, Water 2: 2825±115). Molecular formulae indicative of condensed aromatic structures, of likely pyrogenic origin, were rare in Water 1 and not detected in Water 2 (Supplementary Figure S3, Supplementary Table S7). Most molecular formulae were representative for unsaturated aliphatic, polyphenolic, and highly unsaturated phenolic compounds. The latter comprised on average 97% and 95% of the total signal intensity of each mass spectrum of Water 1 and 2, respectively. In contrast, molecular formulae that are consistent with peptide, fatty acid, and carbohydrate structures were low in relative abundance (<0.05% of overall intensity, <0.3% of formulae count). The dominance of potentially refractory molecules suggested that most DOM in both waters may not be readily degradable by microorganisms on the timescales investigated in this study. Nevertheless, we revealed significant compositional changes in DOM during storage (Water 1: canonical R=0.94, P<0.001; Water 2: canonical R=0.96, P<0.01) that affected a similar group of compounds in both waters (Figure 5, Supplementary Figure S4). These incubation-time associated shifts in DOM composition were also reflected in diversity patterns. Diversity and richness of DOM increased, evenness decreased during storage for Water 1, yet no significant trends were identified for Water 2 (Supplementary Figure S5). The strongest changes in molecular DOM composition occurred at the onset of and during microbial growth, which suggests that these shifts were mainly driven by the growing microorganisms (Figure 5).

**Figure 4.**
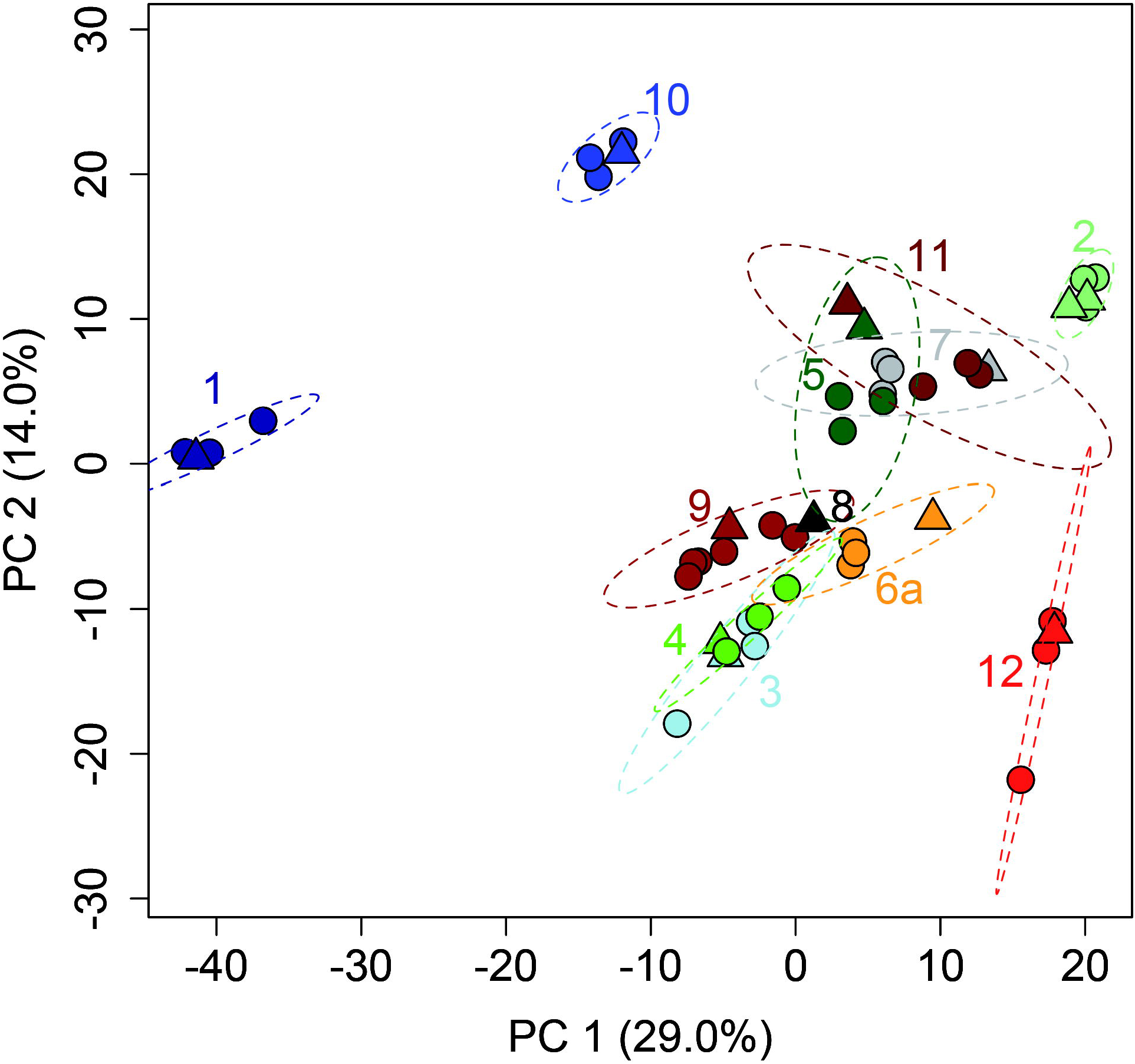
Variation of DOM composition across all well and bottled waters one day after filling. PCA based on log-transformed relative signal intensities (FT-ICR-MS) of identified molecular formulae shows strong compositional contrasts between water types (identical color); the well water samples (triangles) largely cluster with the corresponding bottled water (circles) with some bottle-specific variation. Numbers refer to water types. No data was available for well 6b. Well 8 was described by a single sample. Ellipses correspond to 99% confidence limits of PCA-scores.

**Figure 5.**
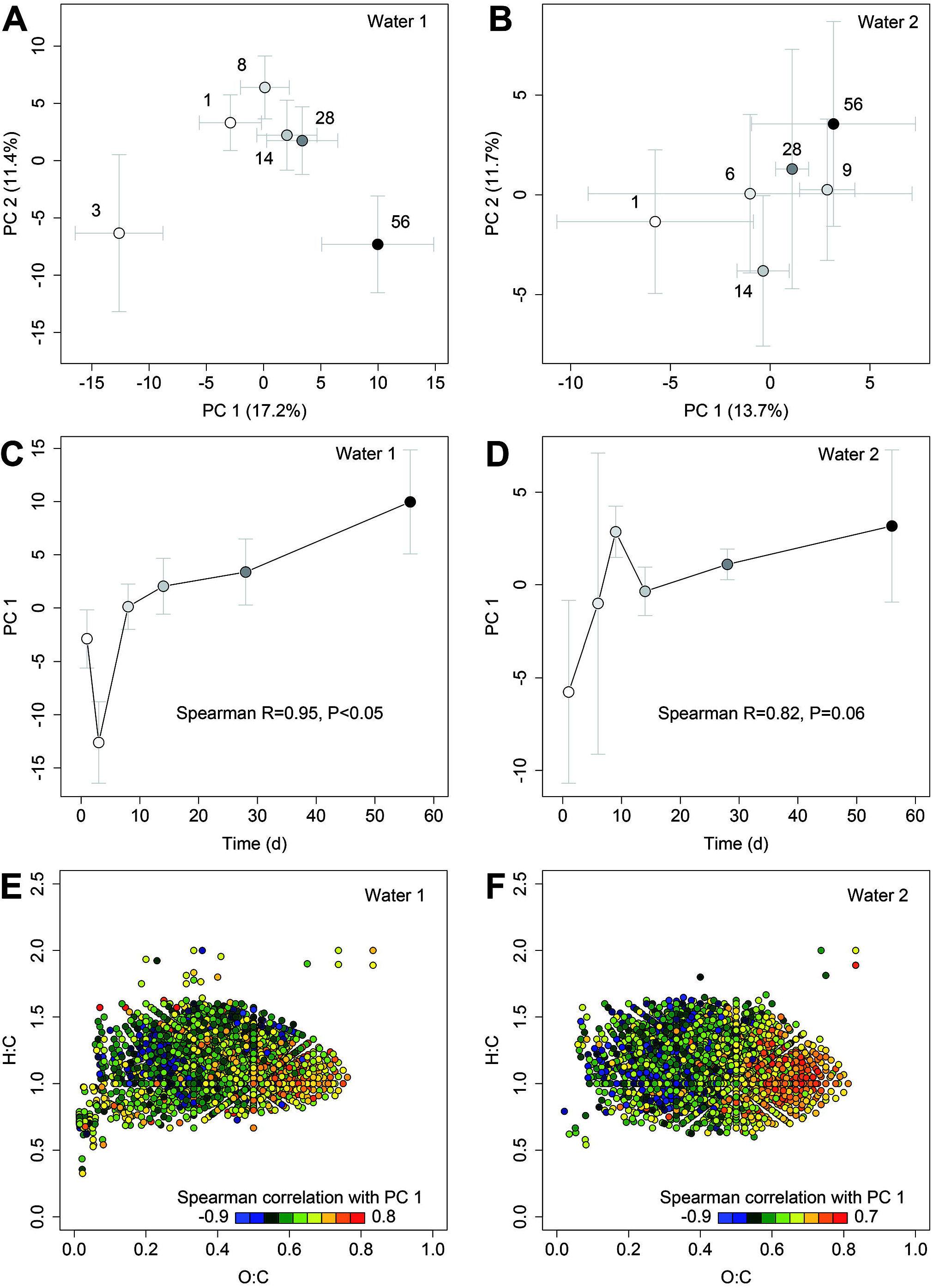
Temporal changes of dissolved organic matter composition in natural mineral water 1 and 2 after bottling. In year 2012, Water 1 and 2 were filled in PET bottles and monitored during 56 days of storage. **(A, B)** PCA based on log-transformed relative signal intensities (FT-ICR-MS) of identified molecular formulae; crosses are mean (±SD) scores of 3 replicates, numbers and coloring refer to incubation time (days after bottling). **(C, D)** Dominant compositional change of DOM (as captured by the PC 1) over incubation time; mean (±SD) scores of 3 replicates. **(E, F)** van Krevelen plots of sum formulae identified by FT-ICR-MS in Water 1 and 2. Each dot represents one sum formula and its location informs about oxygen richness (O:C) and saturation (H:C). Dot color shows Spearman correlation coefficients of relative signal intensity with the PC 1; red (positive correlation) and blue (negative correlation) indicate formulae likely increasing and decreasing over incubation time, respectively. Only sum formulae detected at least 4 times are shown to avoid artificially inflated correlations.

### Core microbiota and diversity of dissolved organic matter in various European bottled waters and corresponding well waters

We further analyzed physicochemical properties (e.g. pH, oxygen), bacterial community composition, and molecular DOM composition in 12 natural mineral waters from Belgium, France, Poland, Switzerland, Spain and the UK (referred to as the ‘diversity study’). As aforementioned, higher oxygen concentrations in freshly bottled water (6.3-18.0 mg/L) than in well waters (4.6-5.7 mg/L) (Table 1, Supplementary Tables S3 and S4) are likely due to pressure filling. Changes in the planktonic and biofilm bacterial communities between day 1 and day 28 of the various bottled water and well water samples were analogous to the time-course experiment. The taxonomic diversity and, based on rarified and resampled datasets, the mean number of observed (44 vs 17), estimated (Chao1: 64 vs 24), and dominant (>1%: 11 vs 5) OTUs were higher at day 1 than at day 28 (Supplementary Figure S6, Supplementary Table S5). Beta-diversity was higher between day 1 samples than between day 28 samples (median resampled p<0.001, two-sample t-test and Mann-Whitney test), which indicates that growth of few, phylogenetically-similar bacteria was driving converging microbiota compositions (Supplementary Figure S7). Only samples from Water/Well 6 contained bacterial communities that differed extensively from other samples at day 28. In general, only between 1 and 15 OTUs (mean 4) collectively contributed >90% of the 16S rRNA gene abundance in each bottled water at day 28 post bottling. At this time point, dominant OTUs in plankton and/or biofilm communities were members of the *Betaproteobacteria* (mostly *Comamonadaceae*, but also *Rhodocyclaceae*, *Oxalobacteraceae*, and *Methylophilaceae*), *Alphaproteobacteria* (*Caulobacteraceae*), *Gammaproteobacteria* (*Nevskiaceae*, unclassified family), and *Spirochaetes* (*Leptospiraceae*) (Supplementary Figures S6 and S8).

Using the combined 16S rRNA sequence dataset from the time-course experiment and the diversity study (215 samples), we determined the core microbiota in three categories of bottled water samples: (1) ‘well water’ samples, (2) ‘early bottled water’ samples from day 1 after bottling that contain the seed microbiota, and (3) ‘late bottled water’ samples from >14 days after bottling that contain the mature microbiota after growth has occurred. We also identified the OTUs that were shared between the core microbiota in these three categories as ‘ubiquitous’ OTUs. Good’s coverages of individual 16S rRNA sequence libraries ranged from 86% to 100% (mean 99%) (Supplementary Table S5), indicating that the majority of OTUs in each sample was detected. We here define the core microbiota, by considering OUT abundance and habitat occupancy [53], as the sum of OTUs that are each present at a relative abundance of ≥0.5% per sample and in ≥50% of all water brands (12 in total) per category.

The sum of all non-redundant OTUs across all samples was 1295, of which only twelve OTUs were identified as core microbiota (Figure 6). These twelve core OTUs belonged to either *Gammaproteobacteria* or *Betaproteobacteria* (Figures 3 and 6). OTU 3 (*Pseudomonadaceae*), OTU 181 (*Oxalobacteriaceae*), and OTUs 1 and 2 (*Comamonadaceae*) were ubiquitous across all three core categories considered. The ‘well water’ core consisted of the four ubiquitous OTUs and OTU 21 (*Enterobacteriaceae*). The shift from ‘well water’ to ‘early bottled water’ was marked by the addition of four additional members (OTUs 4, 17, 1097 and 1183) of the *Gammaproteobacteria*, which were generally more abundant at these early time points (Supplementary Figure S6). The subsequent change from early to late bottled water was characterized by the replacement of four gammaproteobacterial OTUs (OTUs 17, 21, 1097 and 1183) by three *Comamonadaceae* (OTUs 225, 253, 993). At these later time points, the *Comamonadaceae* also dominated the mature microbiota in most waters (Figure 6). Hence, microbial succession during storage of bottled waters was uniquely characterized by a shift from a gammaproteobacterial to a betaproteobacterial core microbiota.

**Figure 6.**
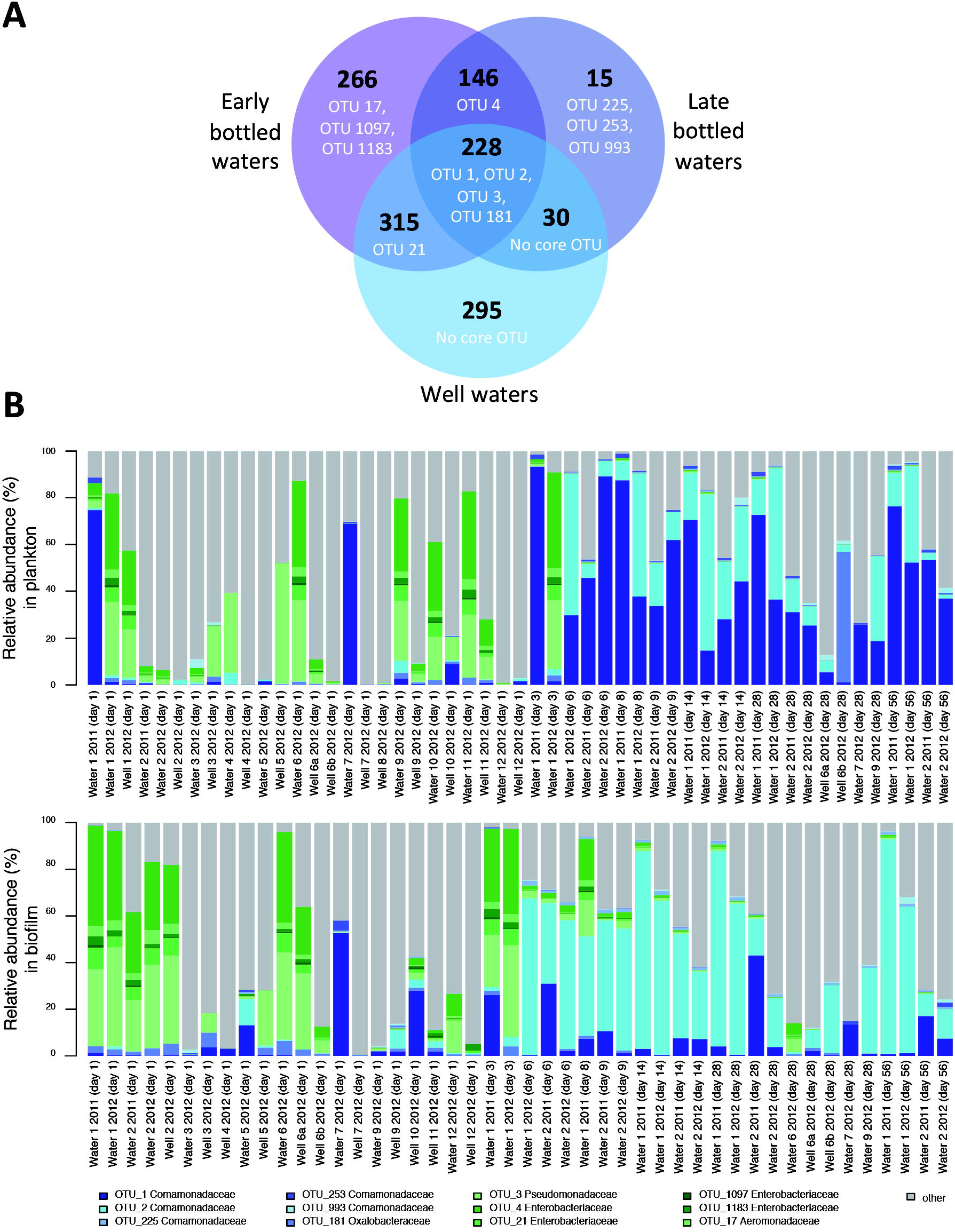
The core microbiota of bottled water. (A) Venn diagram showing core OTUs (in white, with their individual names) and the total number of OTUs (in black) in well waters or bottled waters sampled one day (‘early’) and >14 days (‘late’) after filling in PET bottles and how these are shared between these categories. **(B)** Mean relative abundances of the twelve core OTUs in all analyzed plankton and biofilm samples from different days after bottling. For example, OTU 1 was more abundant in plankton than in biofilm (p = 4.37e-08) and OTU 2 was more abundant in biofilm than in plankton (p = 0.006867) (based on one-sided Wilcoxen Rank Sum test). Furthermore, OTU 1 was more abundant in plankton (p = 0.008156) and less abundant in biofilm (p = 4.423e-05) than OTU 2 (based on paired one-sided Wilcoxen signed rank test).

Early (day 1) bottled and well water samples were analyzed by FT-ICR-MS to characterize and compare their DOM composition and molecular diversity. Collectively, 3813 different molecular formulae were identified across all samples (Supplementary Figure S3). On average approximately 2770 molecular formulae were identified per water sample. Of these, on average 89.8 % are consistent with structures of highly unsaturated phenolics, constituting the most dominant molecular group. Compound groups of potential unsaturated aliphatics and polyphenols accounted for on average 4.9 % each. Together, these three molecular groups covered >99.5% of the total FT-ICR-MS signal intensity. DOM sampled from product-specific bottles largely reflected DOM sampled directly at the wells, yet clear differences in DOM concentration and composition between the various sources were evident (Figure 4) and associated with distinct molecular properties (Supplementary Figure S9). Between-bottle variation was mostly aligned with a minor axis of compositional change (principal component 2) (Figure 4), but was not significantly higher than instrument measurement error, i.e. the compositional variation of a routinely measured natural DOM standard (permutational test for difference in beta-diversities, minimum P=0.51 adjusted by Tukey HSD). For all waters and well samples, DOM consisted of many low-concentrated compounds that are usually not readily available for microbial degradation. This molecular composition with highly diluted individual DOM moieties is consistent with the high estimated age of the source waters (1.5 to 15,000 years) (Supplementary Table S3) [54].

## DISCUSSION

While safe supply of drinking water sustains public health, its microbiological monitoring is traditionally restricted to detecting and managing pathogenic bacteria. A more holistic understanding of drinking water microbial ecology would allow better water management, including predictive assessment of microbiological risks or benefits [55]. Research has largely focused on microorganisms that are detrimental to human health [56] or contribute to the deterioration of water distribution systems [57]. However, the natural aquatic microbiota also provides many beneficial functions for water safety, including resistance against proliferation of pathogens [58, 59] and degradation of contaminants [60]. Technological advances in microbial community and metabolite analysis enable a cohesive assessment of drinking water ecology. Despite its shortcomings e.g. to assess microbiological safety (Supplementary text), multiplexed 16S rRNA gene diversity analysis by next-generation sequencing has revealed insights into the biogeography, assembly, and dynamics of the planktonic and biofilm communities in drinking water distribution systems [53, 61-64] and provided the basis for a first predictive framework for microbiota management [65]. In comparison, knowledge of the microbial ecology of bottled waters, another major drinking water source, is more limited. In contrast to tap water, which is usually used immediately, bottled waters may be stored for several months at ambient temperatures until consumption, conditions in which microorganisms grow rapidly, particularly in non-carbonated natural mineral waters [6, 66].

In this study of natural mineral waters from 12 different European bottled water plants, water obtained directly from wells and freshly bottled, non-carbonated waters both contained a very low number of microbial cells (≤10^4^ cells/mL, Figure 1). These cells can be extremely small, are difficult to cultivate (≤100 colony-forming units/ml), show low metabolic activity, and have low detectability by FISH, likely due to starvation [2, 67-69]. We show that this low-biomass, largely inactive or dormant ‘seed microbiota’ has considerable diversity at various taxonomic levels in each type of bottled water. Not surprisingly, as shown previously for two other bottled water brands [2, 4], bacterial growth and a considerable shift in microbiota composition occurred within about a week after bottling. Here, we demonstrate that only few species from the diverse, low-biomass seed microbiota became metabolically active and grew, resulting in increased similarity of the planktonic and the bottle wall-associated microbiota within and across water types. Furthermore, the timing of shifts in the composition of planktonic and biofilm microbiota during the first two weeks after bottling (Figure 1) indicated that bulk growth first occurred in the aquatic phase. Fewer than 15 OTUs comprised the majority of the community in each bottled water once the microbiota had reached the stationary growth phase. This pattern of community change, which included a characteristic shift from a low-diversity core microbiota of *Gammaproteobacteria* to a low-diversity core microbiota of *Betaproteobacteria* (mostly *Comamonadaceae*), is consistent with a community assembly scenario in which abiotic factors specifically select for growth of functionally and phylogenetically similar and, in this case also, ubiquitous groundwater bacteria [70]. Such environmental filtering or species sorting was recently shown to act during the assembly of microbial communities in rock pools from widespread bacterial taxa [71].

Who are the most important bacteria in bottled water and what are the metabolic features that support their growth? Most species that grew in the bottled waters are related to microorganisms from other oligotrophic groundwater [16] and drinking water ecosystems [53, 61-63] and were also detected in the well waters and immediately after bottling. The core OTUs that were most widespread and grew in different bottled waters (OTUs 1, 2, 225, 253, 993) are all members of the betaproteobacterial family *Comamonadaceae*, including the genera *Curvibacter* (OTUs 1 and 253), *Aquabacterium* (OTUs 2 and 225), and *Polaromonas* (OTU 993) (Figure 3). Members of these genera have been isolated from German drinking water biofilm (*Aquabacterium* species) [52], Japanese well water (*Curvibacter fontanus*) [51], and Swedish drinking water (*Polaromonas aquatica*) [72]. All are mesophilic chemoheterotrophs that grow under aerobic/microaerophilic conditions. Consistent with previous findings [2, 4], these genera are autochthonous microbiota members in bottled natural mineral water and are widely distributed in nutrient-poor drinking waters. OTU 2, with 100% 16S rRNA gene identity to the biofilm-derived *Aquabacterium parvum* type strain (Figure 3), seemed to preferentially colonize the bottle wall as it was significantly more prevalent in biofilm than in plankton samples (Figure 6). Accessory species that grew in bottled water, i.e. non-core OTUs that were abundant after growth in one or only few water types or samples, might either be functionally redundant to the core microbiota members or metabolize substrates that are only present in certain water types or transiently present in certain water samples. Such sporadically-appearing substrates may include single-carbon (C1) compounds as some members of the accessory bottled water microbiota are related to taxa (e.g. *Betaproteobacteria*: *Methylophilaceae*, *Methyloversatilis* in *Rhodocyclaceae*, *Alphaproteobacteria*: *Methylobacteriaceae*) that contain obligate and facultative methano-/methylotrophs [73]. Methano-/methylotrophic taxa are also found in drinking water distribution systems and it has been suggested that methanogenesis in anoxic sites of the source water aquifer provides methylated substrates to fuel their metabolism [53].

What are the physicochemical constraints in the different bottled waters that select for such a highly similar microbiota? All waters were stored in PET bottles at 20-24°C in the dark, and remained well oxygenated and at a circumneutral pH during one to two months of storage. Total DOM concentrations varied between waters but were low (<1.2 mg/L) as in other bottled natural mineral waters [8]. These uniform habitat characteristics among all investigated waters set the basic metabolic conditions for microbial growth. In order to describe the chemical environment for growth of the bottled water microbiota in more detail, we employed ultra-high resolution mass spectrometry to characterize DOM composition in all waters and its changes during bacterial growth in two waters with maximally different DOM concentrations. Diversity of DOM, i.e. the number of identified DOM moieties, was variable between the various types of waters, and, indeed, each water had a characteristic DOM composition. Despite these differences, DOM in all waters was dominated by molecular groups that are apparently unsusceptible to immediate microbial degradation, such as highly unsaturated phenolic compounds. A unifying characteristic of the investigated waters is that they were derived from relatively old subsurface water masses that contain aged, largely refractory, humic-like DOM, similar to DOM from other subsurface aquatic environments such as karst pools [16]. The predominance of polyphenols and highly unsaturated compounds is characteristic for DOM originating from vascular plant debris and thus from terrestrial sources. Additionally, the high molecular diversity is indicative for highly processed soil-derived DOM. In contrast, fresh plant leachates and microbially-derived DOM that are typically enriched in more saturated compounds were not significant components of the bottled water DOM.

Despite lack of peptide and carbohydrate molecular formulae and the predominance of soil-derived DOM, part of the DOM could have served as substrate for the bottled water microbiota. During incubation of Waters 1 and 2, specific molecular formulae indicative of polyphenolic structures decreased or increased in relative abundance (Figure 5EF). Relative decreases could be due to adsorption to the bottle wall or microbial degradation. Degradation of soil-derived polyphenols by the riverine microbial community was shown in a study of the Amazon River [74]. In addition, the numbers of chemically distinct molecules increased over incubation time in Water 1, suggesting an additional contribution of products of microbial metabolism to the DOM pool [75]. Despite notable differences in DOM concentration and composition, compositional change of DOM spanning the time-course experiment were similar for both waters (Figures 5 and 6) and might be reciprocally linked to the growth of only few, related OTUs, which also caused the shift to similar low-diversity bacterial communities. Significant changes in DOM composition were revealed in both waters, even though Water 2 had a much larger background of refractory molecules and lost a much smaller percentage of its overall DOM.

Growth of only few bacterial species was related to the significant turnover of likely hundreds of multi-carbon molecules. Physiological flexibility and simultaneous utilization of multiple substrates, known as ‘mixed substrate growth’ [9], are key adaptive features that allow microorganisms to compete and grow in oligotrophic water environments with only small concentrations of individual DOM moieties. It is tempting to speculate that individual bacterial species growing in bottled waters are substrate generalists and simultaneously forage on multiple, low-concentrated organic compounds in the diverse DOM pool. Additionally, the more refractory molecules in the DOM pool might become an increasingly important nutrient source for bacteria over prolonged storage of bottled water beyond the time frame investigated in this study. This could contribute to continuous compositional changes in the bottled water microbiota [4], maintenance of constantly high cell numbers, and only little decrease in cultivability [5, 6, 66, 76] as observed over several months after bottling.

## CONCLUSIONS

This study presents the most exhaustive view to date of the microbiota in bottled natural mineral water and the interplay of its individual species with their physicochemical environment, including the highly complex DOM matrix. Despite the high molecular diversity of available DOM, the actual microbial niche space in these closed aquatic ecosystems appears rather restricted as demonstrated by the high similarity and low richness of bacteria that grew in different bottled waters. We show that turnover of hundreds of chemical molecules, from a background of >2500 DOM molecules, was related to the growth of less than ten abundant species that were recruited from a seed community of up to a few hundred bacterial species. Although only a small fraction of both the chemical and microbial richness was involved, the physiologically active bacteria seemed to utilize many different and low-concentrated DOM molecules simultaneously [9]. Assembly of a characteristic low-diversity bottled water microbiota after bottling was mainly driven by abiotic factors typical for the bottled water environment. Similar shifts of DOM composition during post-bottling microbial growth suggested that consumed and produced fractions of DOM are rather similar across various waters. The composition of bioavailable DOM may thus act as an important selection factor. Biotic factors, including cross-feeding on products from primary degraders of complex recalcitrant DOM molecules, may become progressively more relevant for community maintenance and dynamics during continuous storage of bottled waters over several months. We show that *Curvibacter*, *Aquabacterium*, and *Polaromonas* (*Comamonadaceae*) are habitat generalists that represent the growing core of mesophilic, heterotrophic, and aerobic bacteria in many oligotrophic natural mineral waters. Complementarily, the composition of habitat specialists, species that grew only occasionally or only in specific bottled water types, defined the ‘microbial uniqueness’ of each bottled water. These findings are in agreement with the perception that drinking waters from different sources could be distinguished by their unique microbial and chemical composition [62]. These new insights into the microbial ecology of bottled natural mineral waters, together with the established 16S rRNA gene sequence and DOM molecular datasets, are an important knowledge base and data resource for water source tracking and for developing new microbial and molecular markers for improved water quality monitoring.

## ETHICS APPROVAL

Not applicable.

## CONSENT FOR PUBLICATION

Not applicable.

## AVAILABILITY OF DATA AND MATERIALS

Sequence data supporting the conclusions of this article is available in the NCBI Sequence Read Archive (SRP091585) and GenBank (KX989480 to KX989499). The raw FT-ICR-MS dataset (as absolute peak intensities) is included within the article as Supplementary Table S8.

## AUTHORS’ CONTRIBUTIONS

CCL, GAS, CG, and AL designed the study; CG and CCL coordinated sample collection; CCL retrieved sequence data; CCL, DB, and CWH analyzed sequence data; CCL and CP performed fluorescence microscopy; CCL, CG, XLC, and SG performed physicochemical analysis; GAS, CCL, TD, and JN performed ultra-high mass spectrometry analysis; GAS, DB, and CWH performed statistical analysis; AL wrote the paper with contributions from CCL, CWH, and GAS; All authors revised and approved the manuscript.

## ACKNOWLEDGMENTS

Not applicable.

